# Changes in muscle strength and moderators of protein turnover in a rodent model of anorexia nervosa and recovery

**DOI:** 10.1101/2025.09.29.679271

**Authors:** Megan E. Rosa-Caldwell, Lauren Breithaupt, Ursula B. Kaiser, Ruqaiza Muhyudin, Seward B. Rutkove

## Abstract

Anorexia nervosa (AN) is a psychiatric disorder characterized by severe caloric restriction, leading to health complications. In addition to fat loss, AN also results in profound skeletal muscle loss, yet molecular pathways underlying these musculoskeletal complications or how long-lasting these musculoskeletal consequences may be are currently unknown. The purpose of this study was to investigate the effects of AN and subsequent weight recovery on muscle strength, size, and moderators of protein turnover in a rat model of AN. Female Sprague Dawley rats (n=11/group, 8 weeks of age) underwent 30 days of simulated AN (50-60% food restriction) followed by varying recovery periods. Muscle mass, strength, and protein synthesis/degradation pathways were assessed. AN led to substantial reductions in muscle mass and strength. While muscle mass recovered within 30 days, muscle strength remained depressed in rats with a prior history of AN, suggesting alterations to muscle quality. Moreover, moderators of protein synthesis (Igf1, Redd1, Deptor) remained altered following 30 days of AN and subsequent recovery. These findings suggest muscle impairments in AN may be longer-lasting than previously thought and may contribute to increased health complications and reduced quality of life in those with a history of AN.

## Introduction

Anorexia nervosa (AN) is psychiatric disorder characterized by fear of weight gain and caloric restriction relative to an individual’s energetic needs^1^. These factors result in profound weight loss, leaving people with this condition severely underweight for height and sex^1^. Prevalence rates are difficult to fully determine due to diagnostic and reporting challenges, but it is estimated ∼4% of all women either actively have AN or have a history of AN^2–4^, accounting for millions of individuals and associated healthcare costs. With this severe energy restriction, there is an increase in health complications and morbidity in this population. A recent report found risk of premature mortality is 3.02 times greater in those with AN (or prior history of AN) compared to those without AN throughout an individual’s lifetime^5^. Specifically, odds of premature death from metabolic disorders are almost 20-fold greater in those with AN compared to healthy controls^5^. Skeletal muscle, by mass, as the largest metabolic organ in the body, has substantial impact on overall metabolic health. However, skeletal muscle physiology has <u>not been thoroughly</u> been investigated in this population.

Along with fat loss during AN, there is a concurrent reduction in skeletal muscle. Clinical data suggest that there is ∼24.5% lower muscle mass and ∼35% lower muscle strength in those with AN compared to healthy controls^6^, though differences in methodologies make an exact estimate of muscle alteration difficult to quantify. Moreover, many of these musculoskeletal derangements appear to remain even after weight restoration, with ∼9% lower muscle mass in those in recovery from AN^6^. However, the different recovery protocols and durations of recovery make it difficult to fully identify the longevity of muscle derangements in clinical samples. Correspondingly, there is little understanding of how AN affects muscle health acutely during the low-weight phase of the disease or for how long these periods of muscle deterioration persist.

Skeletal muscle size and strength are maintained by a delicate balance between protein synthesis and protein degradation. Multiple cellular pathways exist that can augment or reduce protein synthesis/degradation^7, 8^. There is limited knowledge as to what extent alterations in protein synthesis and/or protein degradation underlie changes to muscle size during AN. For example, one study found evidence for ∼55% depressed whole body protein synthesis in human patients^9^. Other studies have found enhanced systematic inflammation in patients with AN, which could contribute to muscle loss during AN^10^. However, overall mechanistic insights into the factors contributing to muscle atrophy during the low-weight stages of AN or following weight restoration interventions are lacking.

In sum, muscle physiology has not been thoroughly investigated in this population, either acutely or with weight restoration, across a range of recovery periods. Therefore, the purpose of this study was to evaluate multiple components of muscle strength, size, and protein synthetic and degradative pathways during simulated AN and subsequent weight recovery in a rodent model. We hypothesized that simulated AN would result in severe alterations to size and strength, which would not fully resolve upon weight recovery, and that these would be associated with persistent alterations in protein synthetic and degradative pathways.

## Methods

### Ethical approval and animal methods

All experiments were approved by Beth Israel Deaconess Medical Center Institutional Animal Care and Use Committee. All rats were housed in a temperature (∼23° C) controlled and 12:12 light-dark cycle animal facility. For this study, experiments were divided into shorter term recovery and longer term recovery. The procedures for inducing simulated AN in these rats as well as recovery have been published previously^11^. Briefly, this model recapitulates many clinical characteristics of AN in human participants, in particular low fat mass, low bone mineral density, altered estrous cycle, and increased activity^11^. The duration of this simulated AN protocol also exceeds other rodent models of AN, such as the Activity-Based-Anorexia (ABA) model, where rats lose 20-25% of bodyweight within 7-10 days^12–16^. For both shorter and longer term recovery experiments, 8-week-old female Sprague Dawley rats underwent 30 days of simulated AN, which included consumption of 50-60% less food than *ad libitum* consumption (measured for three days prior). Following the 30-day simulated AN intervention, rats were divided into either shorter-term or longer-term recovery cohorts. Within the shorter-term recovery cohort, groups included acute AN (no recovery, AN-0R, n=11), AN plus five days recovery (AN-5R, n=11), AN plus 15 days recovery (AN-15R, n=11) or healthy controls (CON-0R, n=11). CON-0R rats were age-matched to AN-0R rats. For the longer-term recovery cohorts, groups included AN plus 30 days recovery (AN-30R, n=11), matching recovery time to duration of food restriction, and healthy controls that were age-matched to AN-30R (CON-30R, n=11). For recovery interventions, rats were provided *ad libitum* access to food^11^. A depiction of experimental groups is depicted in Figure 1.

**Figure 1:**
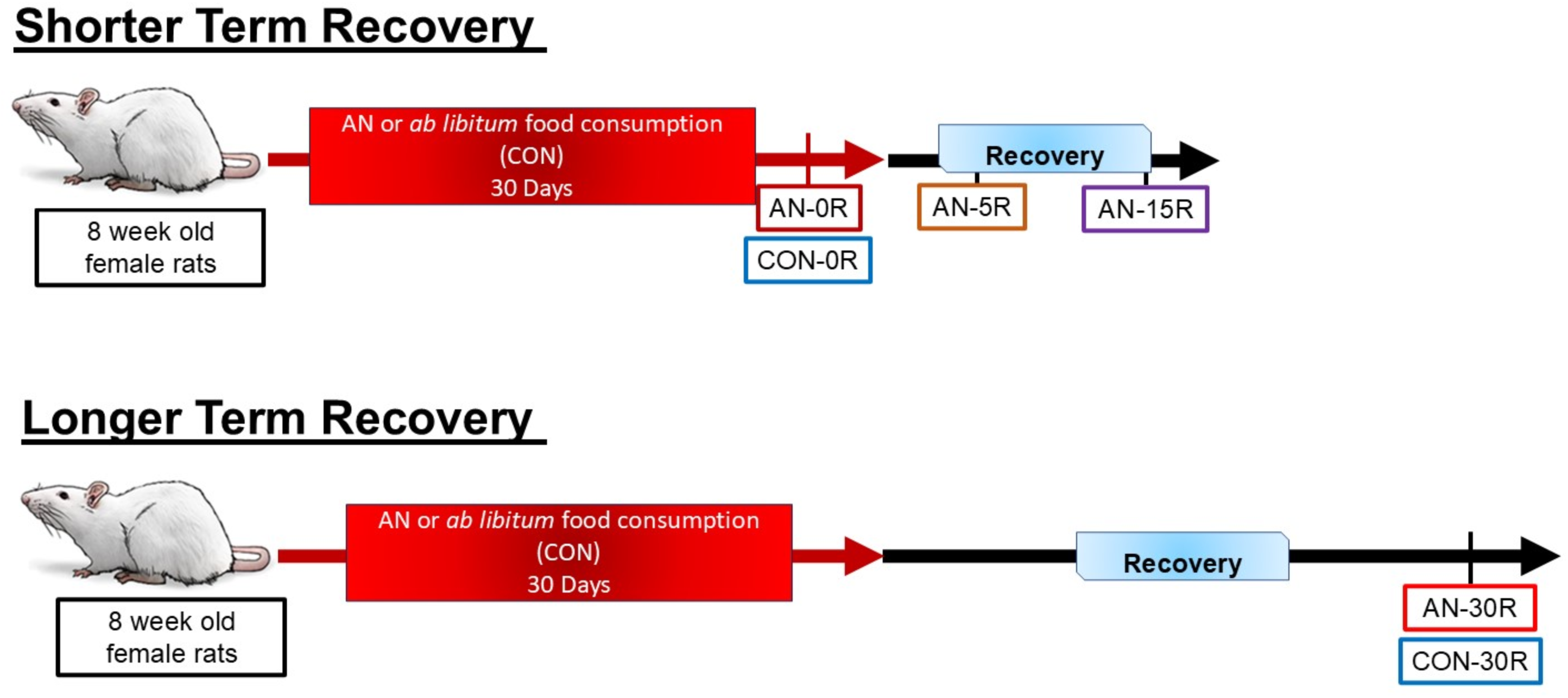
Experimental design for both shorter term recovery and longer term recovery experiments. For shorter term recovery, AN rats underwent 30 days of simulated AN. Then rats were divided into three separate recovery cohorts — no recovery (AN-0R), 5 days of recovery (AN-5R) or 15 days of recovery (AN-15R). Recovery was elicited via reintroduction of ad libitum food consumption. All AN cohorts were compared healthy controls (CON-0R, ad libitum fed) that were age matched to AN-0R rats. For longer term recovery, AN rats underwent 30 days of ad libitum recovery (AN-30R). To account for maturation of rats, AN-30R rats were compared to an age-matched healthy control group (CON-30R) that was fed ad libitum throughout the entire protocol. For all experiments, n=11/group.

After the designated interventions, rats were anesthetized with 2-3% isoflurane, muscle tissues collected and frozen in liquid nitrogen, and rats euthanized by cardiac puncture. ∼20 hours prior to euthanasia, all rats were fasted. One and a half hours (∼90min) prior to euthanasia, all rats were provided with a standard amount of food (3 grams) and given 30 minutes to consume food. This procedure was completed to evaluate acute responses in genes related to protein turnover following an anabolic stimuli (feeding). Additionally, this procedure ensured that feeding status was matched between all rats and there were not significant feeding discrepancies between control rats and various AN rats.

### Volitional strength

Volitional strength was assessed by grip strength, as we have previously described^17, 18^. Briefly, rats’ hindlimb feet were placed on a grip bar attached to a force transducer (Chatillon, Agawam, MA). Rats were then gently pulled from the bar until they released the bar and the corresponding force was recorded. Each rat underwent three trials, with the greatest force being used for analysis. Grip strength was assessed prior to simulated AN and after designated interventions.

### Muscle area, density and fat area, measured via peripheral quantitative computed tomography (pQCT)

Muscle area and density as well as total fat content were evaluated using pQCT, (Stratec, XCT Research SA+, Pforzheim, Germany) as we have previously described^17, 18^. Briefly, rats were anesthetized with 2-3% isoflurane and placed upon a specialized stage with continued anesthesia support. To ensure leg stability, the rat’s leg was place at 0 degrees of knee flexion and the ankle and foot taped to the stage. The rat’s leg was then measured 40mm from the most proximal portion of tibia (found via scout view on the device) with the following parameters: voxel size 0.10 mm, Ct speed 10 mm/sec. Analysis of muscle area, muscle density (a metric of intramuscular fat infiltration), and total fat area was conducted using pQCT software. pQCT scans were completed prior to simulated AN and after designated interventions.

### Maximally stimulated strength

Maximally stimulated dorsiflexion and plantarflexion were completed as we have previously described^17, 18^. Briefly, rats were anesthetized with 2-3% isoflurane and placed on a platform with a footplate force transducer. The rat’s left foot was then taped to the footplate with medical grade tape (3M™ Transpore™ Surgical Tape, Cat# 1527), ensuring no space between the foot and footplate. Needle electrodes were then placed at the knee to elicit either dorsiflexion or plantarflexion. Appropriate needle placement was confirmed with a small (10 Hz) twitch stimulation. Muscles were stimulated at 200 Hz for 200ms to induce maximal tetanus. Tetanus curves were evaluated for maximal force production, time to reach maximal force production, and ½ relaxation time, as previously described^19^. Maximally stimulated dorsiflexion and plantarflexion (measured in that order) were measured prior to simulated AN and after designated interventions. Data were analyzed with baseline values as a covariate to account for possible differences in baseline values.

### Protein turnover

Twenty-four-hour protein turnover was measured as previously described^20, 21^. Approximately 24 hours prior to euthanasia, rats were administered deuterium oxide (D_2_O) via an intraperitoneal injection at a dose of 20 µL/g body weight. Afterwards, rats’ drinking water was replaced with 4% D_2_O in tap water until the following day. Gastrocnemius muscles and plasma were then analyzed for incorporation of deuterium in alanine as a measure of protein synthetic rate, as previously described^22^.

### Histology

The left gastrocnemius muscle was assessed for myosin heavy chain distribution and muscle fiber cross-sectional area^17, 18, 23^. Gastrocnemius muscle was fixed in 10% neutral buffered formalin for 48 hours at 4°C. Afterwards, muscles were rinsed in phosphate buffered saline (PBS). Tissues were mounted in paraffin blocks and sections sliced at 10 µm using a cryotome. Gastrocnemius muscle was then stained for myosin heavy chain I (Cat # ab11083; Abcam), myosin heavy chain II (Cat # ab91506; Abcam), and wheat germ agglutinin (W6748; Thermofisher Scientific). Stained sections were then imaged with a Zeiss Axio Imager M1 Fluorescent Microscope at 20X magnification. Cross-sectional area of muscle fibers was then analyzed with semi-automatic software, as we have previously reported^17, 18, 23^. Each animal had at least 250 fibers analyzed by a researcher who was blinded to all groupings.

### Quantitative Polymerase Chain Reaction (qPCR)

RNA was isolated from gastrocnemius tissue as previously described, using a commercial kit (Purelink RNA Mini Kit, ThermoFisher, Cat #12183025). After ensuring that RNA concentration and quality were sufficient (260/280 ratios >2.1), 1 µg of RNA was then reverse-transcribed to cDNA using commercial kits (Superscript IV VILO, ThermoFisher, Cat #11756500). cDNA was diluted 100-fold and then utilized for qPCR. Quantification of genes of interest were completed using Taqman master mix (Taqman Fast Advance Master Mix, ThermoFisher, Cat # 4444965) and associated probes (Table 1). Gene content was analyzed via ^-ΔΔ^Ct method and were normalized to the housekeeping gene Hprt. Prior to data analysis, we confirmed that Hprt mRNA levels were not different between groups (p>0.05).

**Table 1:** Taqman Probes.

| **Table 1: Taqman Probes** |  |
| --- | --- |
| **Gene** | **Probe Number** |
| Hprt | Rn01527840_m1 |
| Il6 | Rn07315963_g1 |
| Tnfa | Mm00446003_m1 |
| Myostatin | Rn00569683_m1 |
| Atrogin | Rn00591730_m1 |
| MurF1 | Rn00590197_m1 |
| Gadd45a | Rn01425130_g1 |
| Lc3 | Rn02132764_s1 |
| P62 | Rn00709977_m1 |
| Igf1 | Rn00710306_m1 |
| Pax7 | Rn01518732_m1 |
| MyoG | Rn00567418_m1 |
| MyoD | Rn00598571_m1 |
| Deptor | Rn01504430_m1 |
| Redd1 | Rn01433735_g1 |

### Statistical Analysis

Data were divided into short-term recovery (CON, AN, AN-5R, AN-15R) and long-term recovery (CON-30R and AN-30R). For short-term recovery, data were analyzed via one-way analysis of variance (ANOVA). To compare differences between CON and each experimental group, we used Dunnett’s post-hoc adjusted p-value. For long-term recovery, data were analyzed by independent T-test. For all analyses, significance was denoted at p<0.05. To account for variability in baseline values when data was collected longitudinally (e.g., volitional strength, muscle size/density, maximally stimulated force production), data were analyzed with baseline values as a covariate. For cross-sectional measurements, to account for slightly different sizes between rats, we used baseline bodyweight as a covariate. All data were analyzed with SAS (SAS® Studio Release: 3.81, SAS Institute Inc. Cary, NC). All statistical code for this study as well as statistical outputs are available at our Open Science Framework page for this project at https://osf.io/r3t8n/?view_only=32da2b38a51e4742bab2353b635e932a

## Results

### Simulated AN results in loss of bodyweight, muscle mass, and grip strength, which are not fully resolved following weight regain

AN-0R resulted in ∼30% lower bodyweight compared to CON-0R (p<0.001); upon refeeding, AN-5R still had 12% lower bodyweight (Table 2, p<0.001). However, by 15 days of recovery (AN-15R), bodyweight had exceeded CON-0R by ∼5% (Table 2, p=0.042). Some weights of CON-0R and AN-0R rats have been reported previously for these rats^11^. Overall, AN-0R rats had 22-45% (p<0.001) lower muscle weight than CON-0R, depending on the specific muscle, with soleus having the greatest loss in AN rats (∼45%, p<0.001) and plantaris having the least loss (∼22%, p<0.001, Table 2). Five days of recovery (AN-5R) was not sufficient to restore muscle mass, with ∼22% lower gastrocnemius mass (p<0.001), ∼20% lower plantaris mass (p<0.001), ∼16% lower soleus mass (p<0.001), ∼13% lower tibialis anterior mass (TA, p<0.001), and ∼15% lower extensor digitorum longus mass (EDL, p<0.001) compared to CON-0R (Table 2). However, by 15 days of recovery, AN-15R muscle masses were no longer smaller than CON-0R (Table 2, p value range=0.3661-0.999). Of note, the TA mass of AN-15R was ∼9% larger compared to CON-0R (Table 2, p=0.003). AN-0R resulted in ∼20% reduction in rear paw maximal grip strength (Table 2, p=0.017), which was no longer different than CON-0R after 5 or 15 days of recovery (AN-5R (p=0.136), AN-15R (p=0.485), Table 2).

**Table 2:** Muscle weights, Grip Strength, and Food Consumption with simulated AN and short-term recovery.

|  | Group |  |  |  |
| --- | --- | --- | --- | --- |
|  | CON-0R | AN-0R | AN-5R | AN-15R |
| Bodyweight-Baseline (g) | 174.72±3.52 | 172.18±3.52 | 186.45±3.52 | 183.79±3.52 |
| Bodyweight-Final (g) | 216.13±3.06 | 152.02±3.14* | 189.99 ±3.15* | 227.25±3.06* |
| Gastrocnemius (g) | 1.125±0.018 | 0.829±0.019* | 0.872±0.019* | 1.1234±0.018 |
| Plantaris (g) | 0.232±0.005 | 0.181±0.005* | 0.185±0.005* | 0.238±0.005 |
| Soleus (g) | 0.123±0.002 | 0.089±0.002* | 0.104±0.002* | 0.118±0.002 |
| Tibialis Anterior (TA, g) | 0.423±0.007 | 0.333±0.007* | 0.369±0.007 | 0.460±0.007 |
| Extensor Digitorum Longus (EDL, g) | 0.107±0.002 | 0.078±0.002* | 0.090±0.002* | 0.111±0.002 |
| Rear Paw Grip Strength (N) | 6.99±0.31 | 5.70±0.32* | 6.11±0.31 | 6.45±0.31 |
| Overall Average Food Consumption (g) | 14.04±0.08 | 5.97±0.08* | 7.60±0.09* | 9.73±0.08* |
| Average Food Consumption During simulated AN (g) | 14.04±0.07 | 5.97±0.07* | 5.95±0.07* | 6.05±0.07* |
| Average Food Consumption During Recovery (g) | 14.04±0.13 | 5.97±0.13* | 19.81±0.15* | 17.96±0.13* |

Data are depicted as Mean + SEM, n=10-11/group, \* indicates difference between group and CON-0R at Dunnet adjusted $p < 0.05$ . CON-0R and AN-0R had no recovery; therefore their data for Overall Food Consumption, Food Consumption during simulated AN, and Food Consumption during Recovery are the same. Data are corrected with covariate of baseline bodyweight. CON-0R=healthy control age-matched to AN-0R, AN-0R=simulated anorexia nervosa, AN-5R=simulated anorexia nervosa and five days of recovery, AN-15R=simulated anorexia nervosa and fifteen days of recovery.

With longer term recovery, AN-30R still had ∼4% lower bodyweight compared to CON-30R (Table 3, p=0.019). Additionally, AN-30R rats had ∼9% lower gastrocnemius mass (p=0.014), ∼11% lower plantaris mass (p=0.028), ∼13% lower soleus mass (p=0.005), and ∼9% lower TA mass (p=0.028) compared to CON-30R (Table 3). Additionally, AN-30R rats had ∼14% lower rear paw grip strength compared to CON-30R (Table 3, p<0.001).

**Table 3:** Muscle weights, Grip Strength, and Food Consumption with simulated AN and long-term recovery.

|  | CON-30R | AN-30R |
| --- | --- | --- |
| Bodyweight-Baseline (g) | 175.88±3.35 | 181.89±3.35 |
| Bodyweight (g) | 234.58± 2.61 | 224.90±2.61* |
| Gastrocnemius (g) | 1.157±0.026 | 1.053±0.026* |
| Plantaris (g) | 0.247±0.007 | 0.220 ±0.007 |
| Soleus (g) | 0.138±0.003 | 0.120±0.003* |
| Tibialis Anterior (TA, g) | 0.483±0.013 | 0.439±0.012* |
| Extensor Digitorum Longus (EDL, g) | 0.114±0.002 | 0.106±0.003 |
| Rear paw grip strength (N) | 8.47±0.19 | 7.31±0.19* |
| Overall Average Food Consumption (g) | 13.36±0.11 | 10.28±0.11* |
| Average Food Consumption During simulated AN (g) | 13.62±0.13 | 6.08±0.13* |
| Average Food Consumption During Recovery (g) | 13.06±0.09 | 14.67±0.09* |

*Data are depicted as Mean + SEM, n=10-11/group, \* indicates difference between groups at p<0.05. Data are corrected with covariate of baseline bodyweight. CON-30R=healthy control age-matched to AN-30R, AN-30R=simulated anorexia nervosa followed by 30 days of recovery.*

### Simulated AN altered electrophysiological properties of muscle force production, many alterations are not resolved with weight gain

AN-0R rats had ∼21% lower maximal plantar flexion force production compared to CON (Figure 2A, p<0.001). Even after 5 or 15 days of recovery, maximal plantar flexion remained ∼23% and ∼15% lower compared to CON-0R (Figure 2A, p<0.001 and p=0.015 respectively). Time to peak force production (TTP), was ∼15% greater in AN-0R compared to CON-0R and ∼16% greater in AN-5R (Figure 2B, p=0.0178 and p=0.010 respectively). By 15 days of recovery, TTP was no longer different, comparing AN-15R and CON (Figure 2B, p=0.935). Relaxation half-time for plantar flexion was ∼12% greater in AN-0R compared to CON-0R (p=0.032); however, after 5 or 15 days of recovery/weight gain, these differences were no longer significantly different than CON-0R (Figure 2C, p=0.060 and p=0.121 respectively). In analyses of dorsiflexion, although the overall F-test was significant (p=0.039), no differences between CON-0R and each group (AN-0R, AN-5R, AN-15R) reached significance (Figure 2D, p=0.145, p=0.991, and p=0.577 respectively). Similarly, neither TTP nor relaxation half-time were different across groups (Figure 2E and 2F, F-test p=0.080 and p=0.194 respectively).

**Figure 2:**
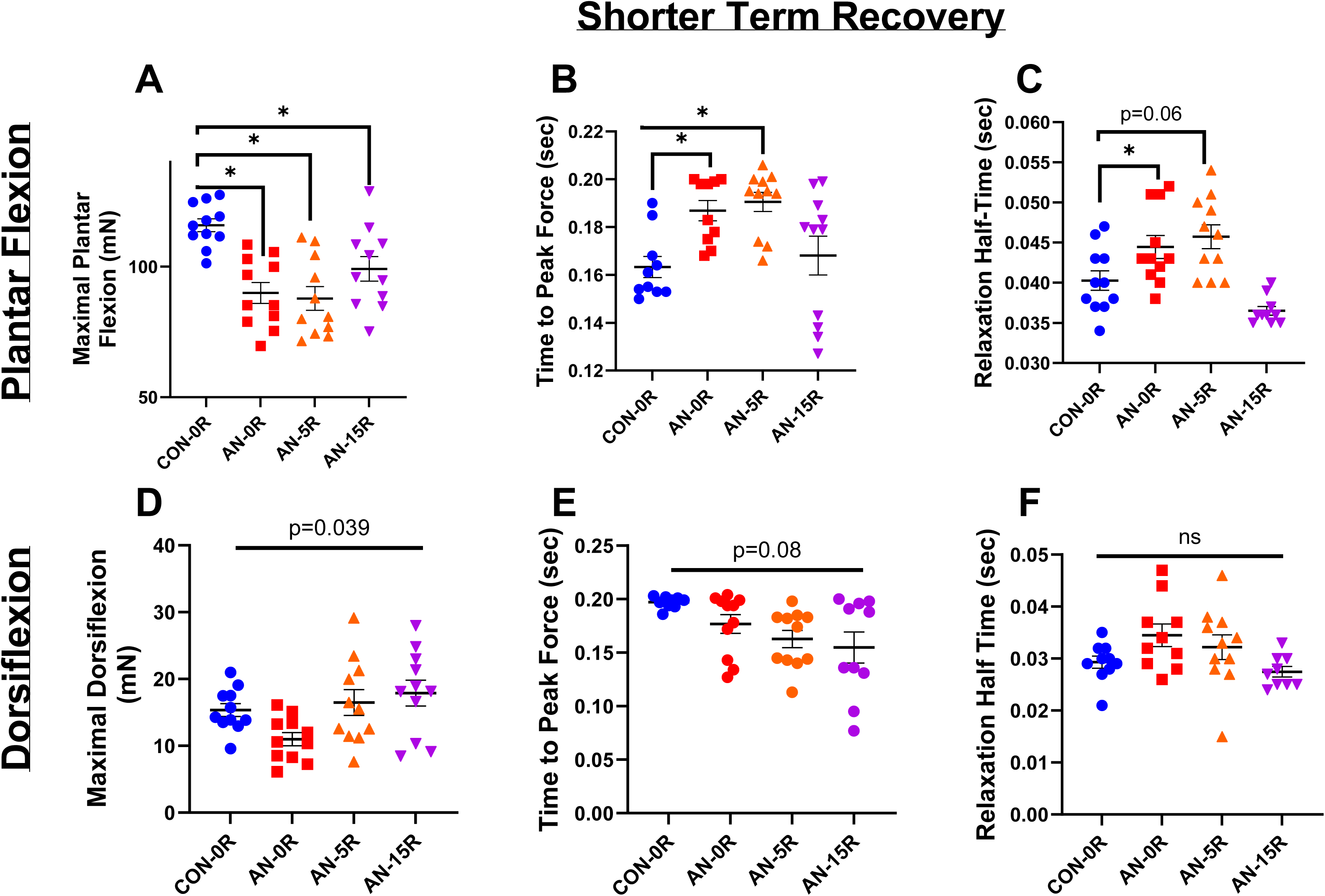
Force production data from simulated AN and shorter term recovery. A) Maximal plantar flexion strength. B) Time to peak force production during maximal plantar flexion. C) Relaxation half-time following maximal plantar flexion. D) Maximal dorsiflexion strength. E) Time to peak force production during maximal dorsiflexion. F) Relaxation half-time following maximal dorsiflexion. CON-0R=healthy controls, AN-0R=simulated anorexia nervosa, AN-5R=simulated anorexia nervosa followed by 5 days of weight recovery, AN-15R=simulated anorexia nervosa followed by 15 days of weight recovery. Data are depicted as mean ± SEM, *=p<0.05. mN*m=millinewtons*meter torque production, Sec=seconds.

With longer recovery, there was still ∼6% lower maximal plantar flexion strength in AN-30R compared to CON-30R (Figure 3A, p=0.021). However, there were no differences between groups in either TTP or relaxation half-time (Figure 3B and 3C, p= 0.885 and p=0.554 respectively). With dorsiflexion, AN-30R had ∼16% lower maximal dorsiflexion strength compared to CON-30R, which approached but did not reach significance (Figure 3D, p=0.073). Similarly to maximal plantar flexion, there were no differences between AN-30R and CON-30R in either TTP or relaxation half-time for dorsiflexion (Figure 3E and 3F, p=0.757 and p=0.848 respectively).

**Figure 3:**
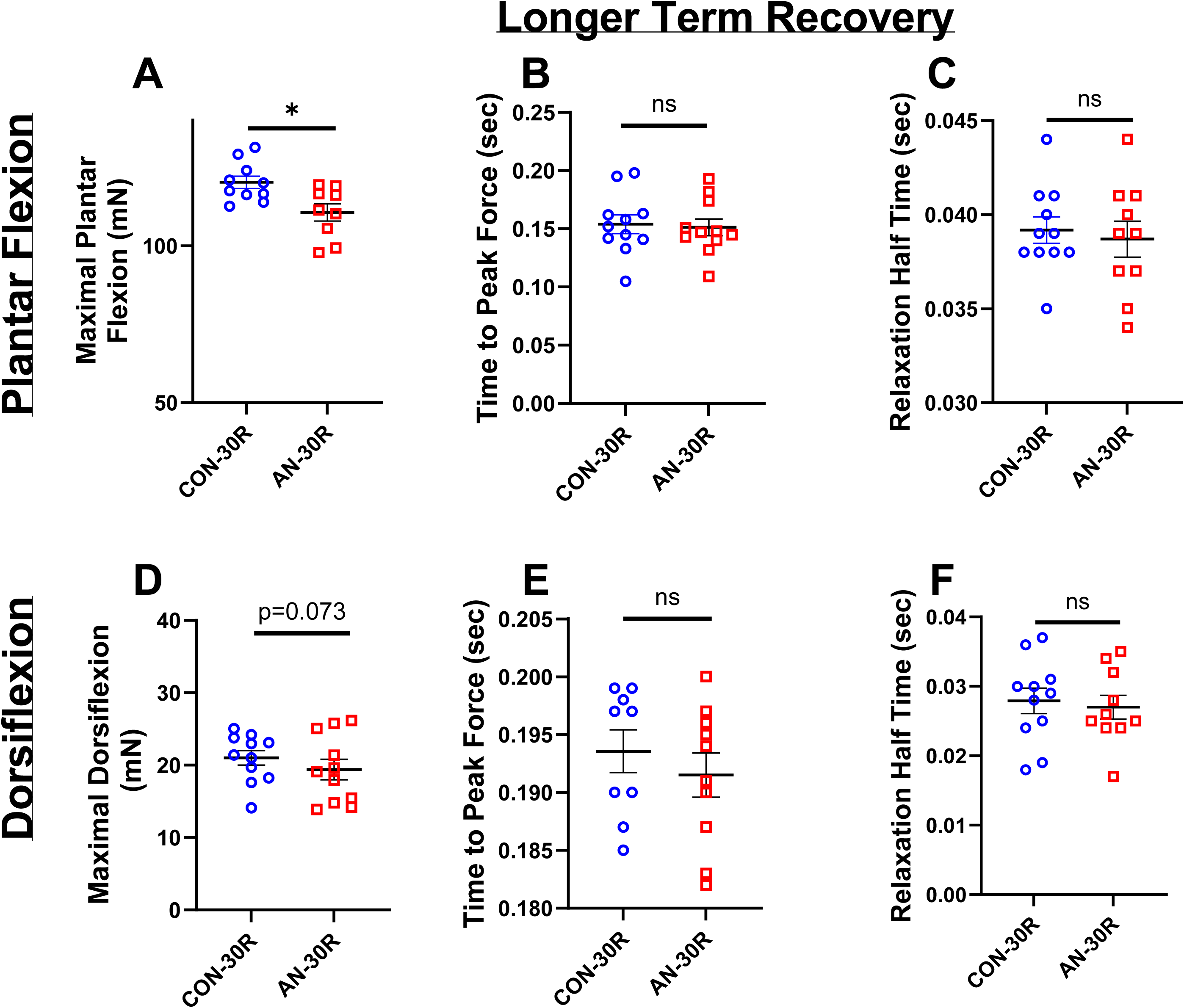
Force production data from simulated AN and control cohorts following longer term (30 days) recovery. A) Maximal plantar flexion strength. B) Time to peak force production during maximal plantar flexion. C) Relaxation half-time following maximal plantar flexion. D) Maximal dorsiflexion strength. E) Time to peak force production during maximal dorsiflexion. F) Relaxation half-time following maximal dorsiflexion. CON-30R=healthy controls age-matched to AN-30R. AN-30R=simulated anorexia nervosa followed by 30 days of weight recovery. Data are depicted as mean ± SEM, *=p<0.05. mN*m=millinewtons*meter torque production, Sec=seconds.

### Simulated AN resulted in loss of muscle and fat, which appears to resolve with weight gain

The AN-0R cohort had ∼23% lower leg (primarily gastrocnemius) muscle area compared to CON-0R as assessed by pQCT (Figure 4A, p<0.001). With 5 days of recovery, AN-5R rats still had ∼17% lower muscle area compared to CON-0R (Figure 4A, p<0.001), yet after 15 days of recovery, muscle area of AN-15R rats exceeded that of CON-0R by ∼11% (Figure 4A, p=0.002). Similarly, AN-0R rats had ∼35% lower fat area compared to CON-0R (p<0.001), which remained ∼23% lower than CON in AN-5R rats (Figure 4B, p<0.001). However, by 15 days of recovery, AN-15R rats had ∼24% greater fat content compared to CON (Figure 4B, p<0.001). Muscle density, a surrogate of intra-muscular fat accumulation, was ∼1 g/mm^3^ greater in AN compared to CON (Figure 4C, p=0.035). By five or 15 days of recovery, AN-5R and AN-15R rats no longer had different muscle density compared to CON (Figure 4C, p=0.279 and p=0.999 respectively). With a longer duration of recovery, AN-30R had similar muscle area and fat area compared to CON-30R (Figures 4D and 4E, p=0.453 and p=0.463 respectively). However, muscle density was ∼1% lower in AN-30R, though this difference did not reach statistical significance (Figure 4F, p=0.106).

**Figure 4:**
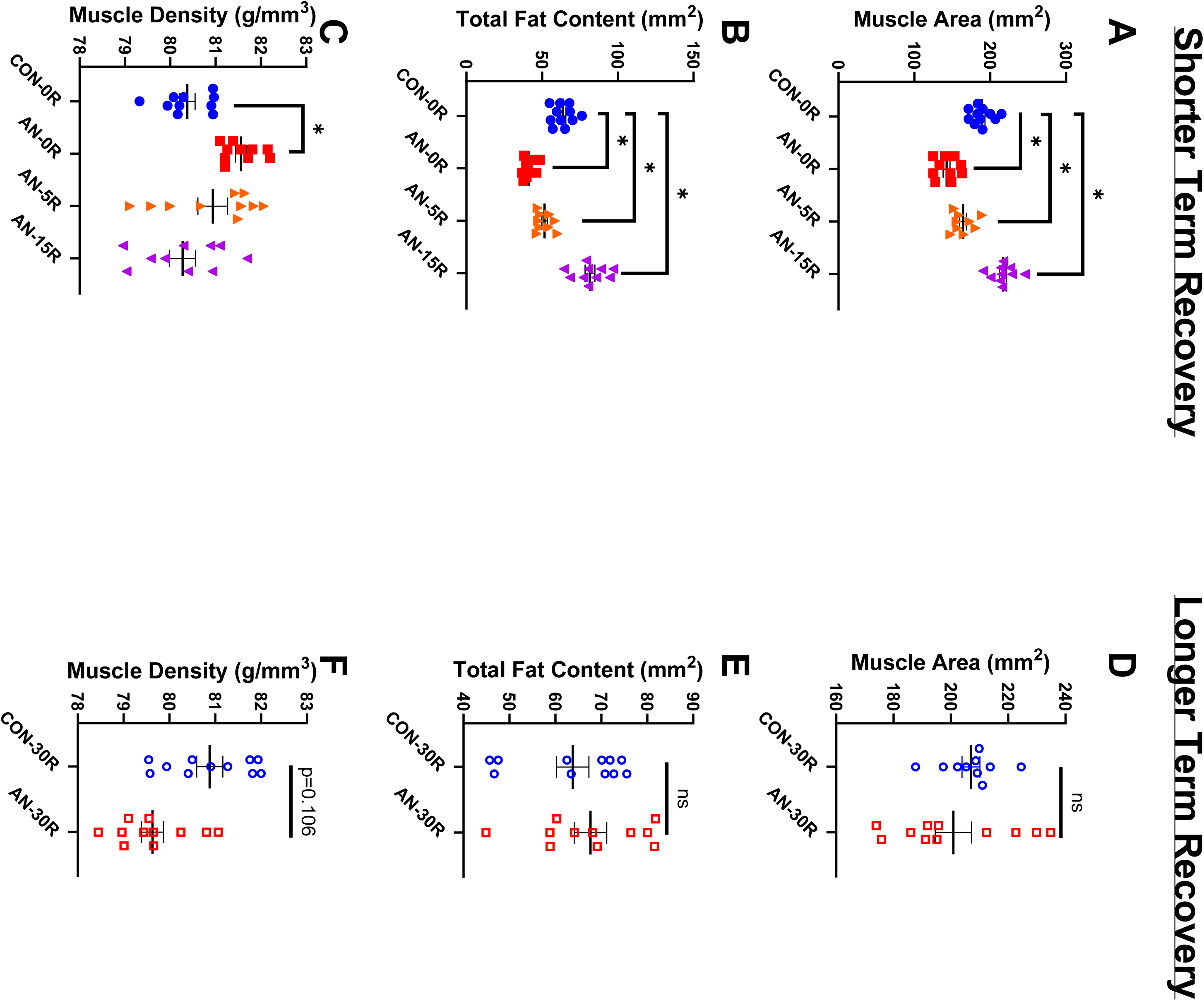
pQCT data from simulated AN and shorter and longer term recovery groups. A) Muscle area during simulated AN and short-term recovery. B) Total fat area during simulated AN and short-term recovery. C) Muscle density during simulated AN and short-term recovery. D) Muscle area data after simulated AN and long-term recovery. E) Total fat area after simulated AN and long-term recovery. F) Muscle density after simulated AN and long-term recovery. CON-0R=healthy controls, AN-0R=simulated anorexia nervosa, AN-5R= simulated anorexia nervosa followed by 5 days of weight recovery, AN-15R=simulated anorexia nervosa followed by 15 days of weight recovery. CON-30R=healthy controls age-matched to AN-30R. AN-30R=simulated anorexia nervosa followed by 30 days of weight recovery. Data are depicted as mean ± SEM, *=p<0.05.

### Simulated AN resulted in reduced muscle fiber cross-sectional area, which was associated with changes in Myosin Heavy Chain Type I fibers

AN-0R rats had significantly lower (29% lower, p<0.001) muscle fiber cross-sectional area (CSA) compared to CON-0R, which was still ∼13% lower in AN-5R compared to CON-0R, although this difference only approached significance (Figure 5A, p=0.078). By 15 days of recovery, fiber CSA was no longer different than CON-0R (Figure 5A, p=0.942). Within fibers positive for myosin heavy chain I (MyHC I), AN-0R rats had ∼26% lower fiber CSA (p<0.001), which remained ∼16% lower in AN-5R compared to CON-0R (Figure 5C, p=0.025), but were not different between AN-15R and CON-0R (p=0.999).There were no differences between any groups in CSA of fibers expressing myosin heavy chain II (MyHC II, Figure 5D, global F p=0.292). In fibers expressing both MyHC I and MyHC II (hybrid fibers), AN-0R rats had ∼29% lower CSA compared to CON (p=0.010), but these differences were resolved in AN-5R and AN-15R (Figure 5E, p=0.145 and p=0.506 respectively). Following a longer duration of recovery, there were no differences between AN-30R and CON-30R in muscle fiber CSA with either MyHC I or MyHC II expression, or both (Figure 6A-E p=0.842, p=0.720, p=0.518, and p=0.685 respectively). There were no differences across any groups in the relative percentages of different MyHC isoforms (data not shown).

**Figure 5:**
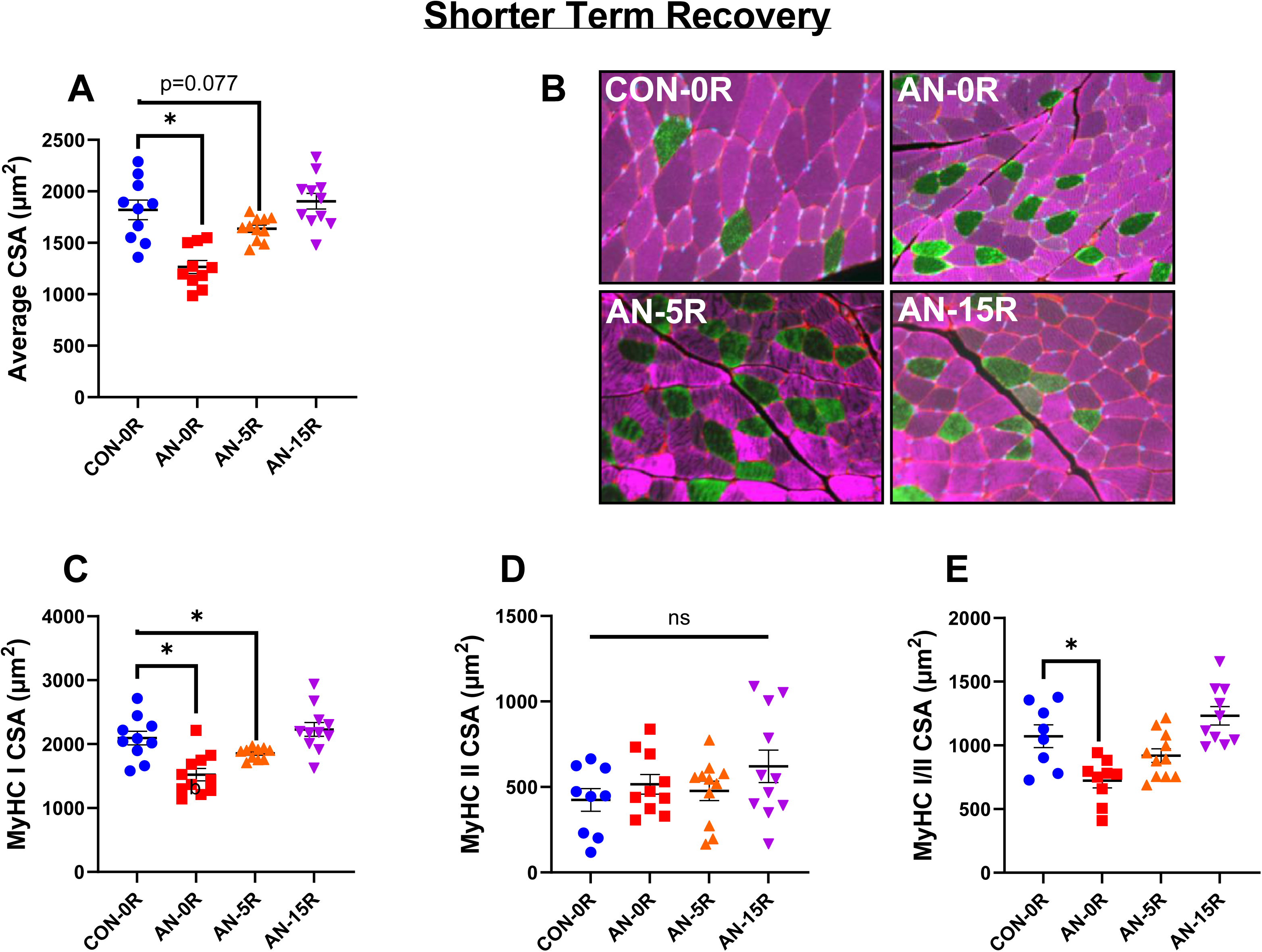
Muscle fiber data from simulated AN and shorter term recovery. A) Mean fiber cross-sectional area (CSA). B) Representative images, collected at 20X magnification. Pink=Myosin Heavy Chain I (MyHC I), Green=Myosin Heavy Chain II (MyHC II). C) Fiber area for MyHC I fibers. D) Fiber area for MyHC II fibers. E) Fiber area for fibers expressing both MyHC I and MyHC II. CON-0R=healthy controls, AN-0R=simulated anorexia nervosa, AN-5R=simulated anorexia nervosa followed by 5 days of weight recovery, AN-15R=simulated anorexia nervosa followed by 15 days of weight recovery. Data are depicted as mean ± SEM, *=p<0.05.

**Figure 6:**
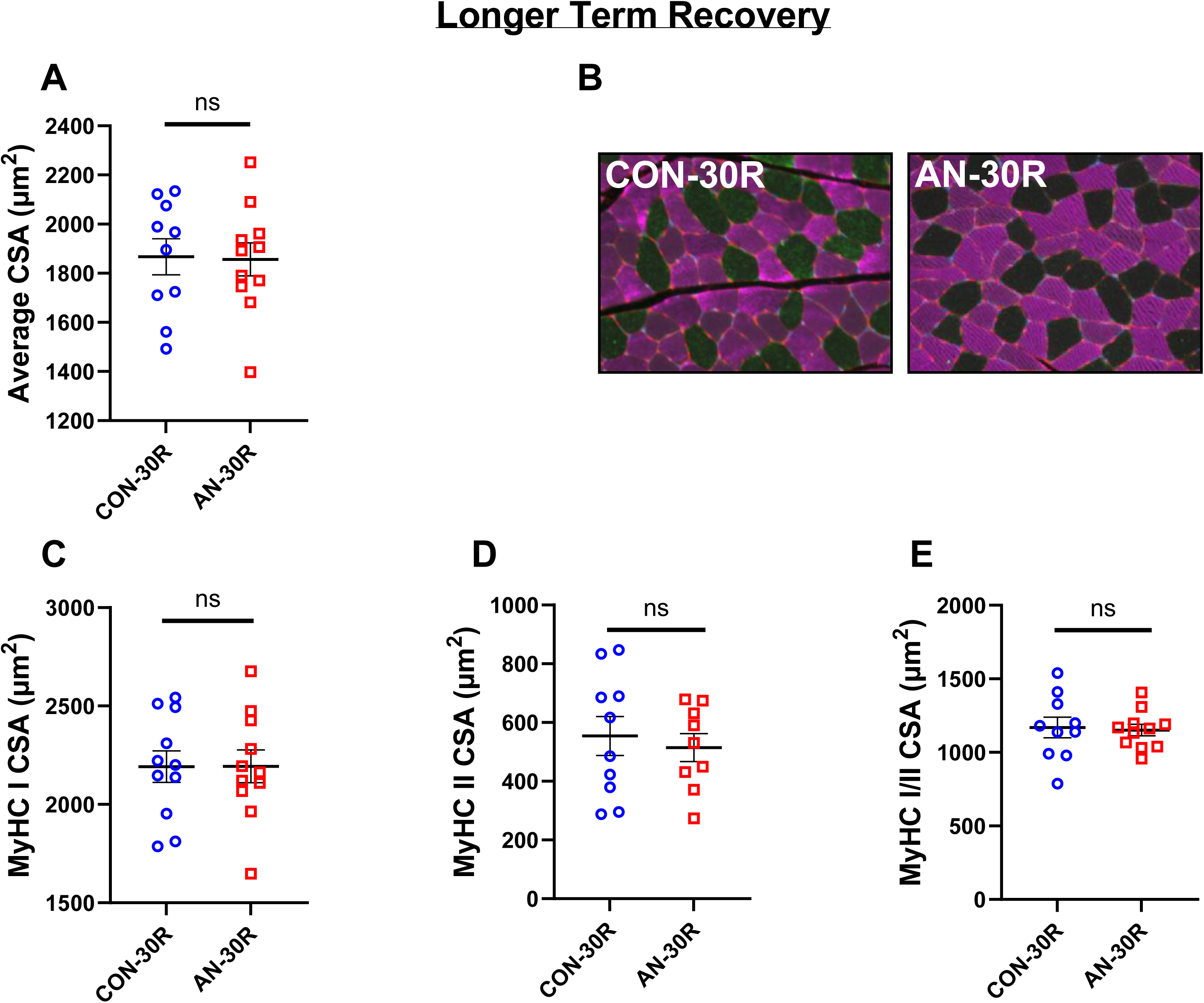
Muscle fiber data following simulated AN and longer term recovery. A) Mean fiber cross-sectional area (CSA). B) Representative images, collected at 20X magnification. Pink=Myosin Heavy Chain I (MyHC I), Green=Myosin Heavy Chain II (MyHC II). C) Fiber area for MyHC I fibers. D) Fiber area for MyHC II fibers. E) Fiber area for fibers expressing both MyHC I and MyHC II. CON-30R=healthy controls age-matched to AN-30R. AN-30R=simulated anorexia nervosa followed by 30 days of weight recovery. Data are depicted as mean ± SEM, *=p<0.05.

### Moderators of protein synthesis were altered with AN, many of which remain altered after weight restoration

The 24-hour fractional synthesis rate (FSR) approached but did not reach significance (p=0.103) between CON-0R and any AN groups (Figure 7A). Anabolic Igf1 mRNA content was not different between AN-0R and CON-0R groups (p=0.978) but was ∼82% and ∼107% greater in AN-5R and AN-15R, respectively, compared to CON-0R (Figure 7B, p=0.015 and p<0.001 respectively). There were no differences in satellite cell marker Pax7 mRNA content between CON-0R and any AN group (Figure 7B, p=0.130). The ANOVA F-test for myogenesis marker MyoG mRNA approached significance (p=0.066), but there were no statistical differences between groups (Figure 7B). Muscle differentiation marker MyoD mRNA content was ∼60% lower in AN-0R and ∼65% lower in AN-15R, compared to CON-0R (Figure 7B, p=0.006 and p=0.002 respectively). Inhibitor of protein synthesis, Deptor mRNA content was ∼40% lower in AN-0R compared to CON-0R, although this difference did not reach statistical significance (p=0.100, Figure 7C). Another inhibitor of protein synthesis, Redd1 mRNA content was ∼50% lower in AN-0R (p=0.022) and remained ∼30% lower (p=0.155, NS) and ∼55% lower (p=0.006) in AN-5R and AN-15R, respectively (Figure 7C).

**Figure 7:**
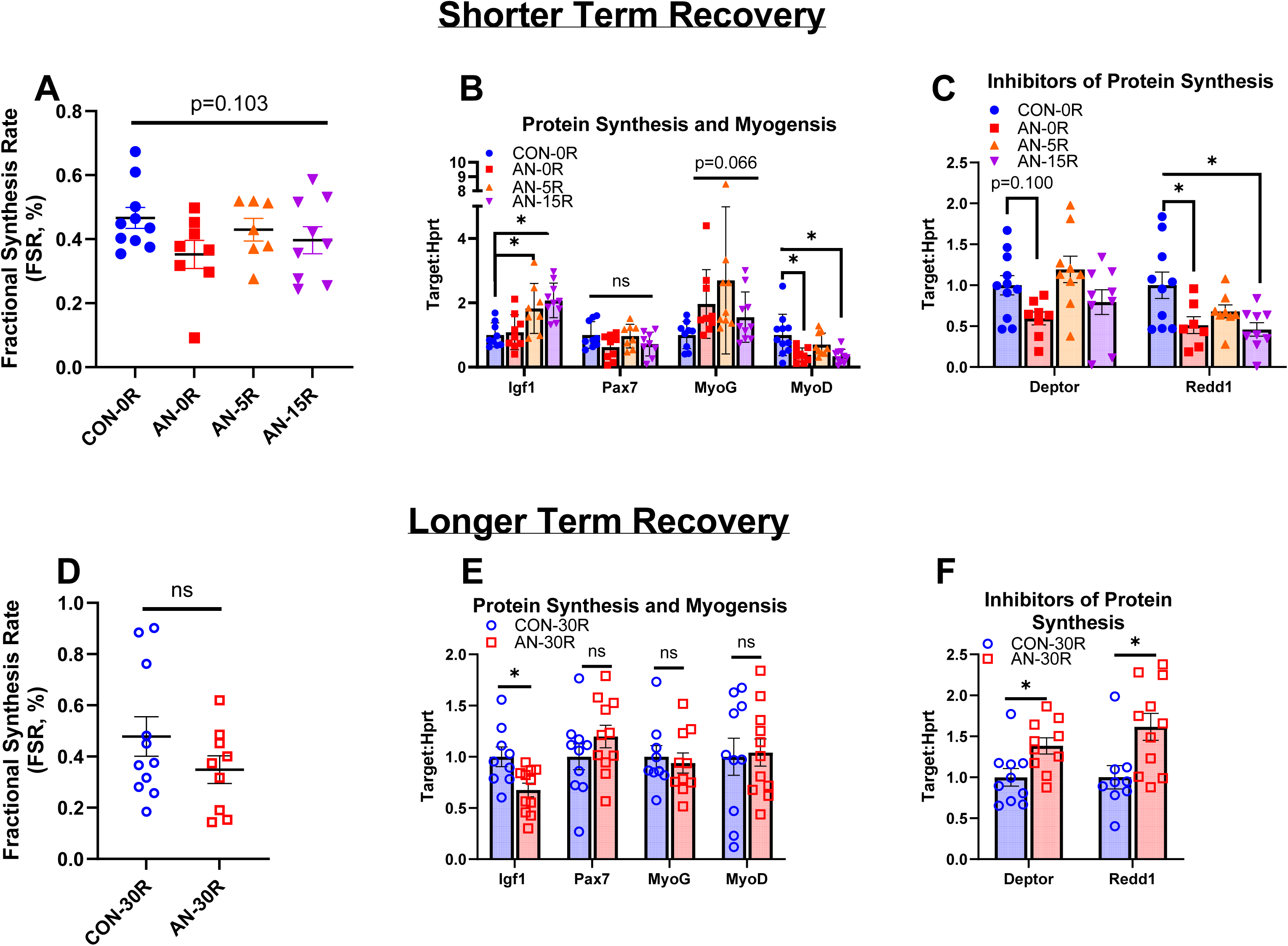
Protein synthesis rate and mRNA data for mediators of myogenesis and protein synthesis. A) Fractional protein synthesis rates after simulated AN and shorter term recovery. B) mRNA content of genes related to protein synthesis and myogenesis after simulated AN and shorter term recovery. C) mRNA content of genes related to inhibition of protein synthesis after simulated AN and shorter term recovery. D) Fractional protein synthesis rates following simulated AN and longer term recovery. E) mRNA content of genes related to protein synthesis and myogenesis following simulated AN and longer term recovery. F) mRNA content of genes related to inhibition of protein synthesis following simulated AN and longer term recovery. CON-0R=healthy controls, AN-0R=simulated anorexia nervosa, AN-5R=simulated anorexia nervosa followed by 5 days of weight recovery, AN-15R=simulated anorexia nervosa followed by 15 days of weight recovery. CON-30R=healthy controls age-matched to AN-30R. AN-30R=simulated anorexia nervosa followed by 30 days of weight recovery. Data are depicted as mean ± SEM, *=p<0.05.

With longer term recovery, FSR was not different between AN-30R and CON-30R (Figure 7D, p=0.676). Igf1 mRNA was ∼33% lower in AN-30R compared to CON-30R (p=0.011), whereas there were no differences in Pax7, MyoG, or MyoD mRNA content between AN-30R and CON-30R (Figure 7E, p=0.249, p=0.681, and p=0.854 respectively). Lastly, Deptor and Redd1 mRNA content was ∼40% greater and ∼60% greater respectively in AN-30R compared CON-30R (Figure 7F, p=0.017 and p=0.013 respectively).

### Inflammatory markers were not different with AN; however, some moderators of autophagy were different with AN and with short-term recovery

There were no differences in muscle inflammatory markers Il6 or Tnfα mRNA content between CON-0R and any AN recovery groups (Figure 8A, p=0.319 and p=0.166 respectively). Muscle specific inflammatory marker myostatin mRNA was ∼126% greater in AN-0R compared to CON-0R (p=0.008), but no differences were noted in AN-5R or AN-15R compared to CON (Figure 8A, p=0.248 and p=0.722 respectively). There were no differences across groups in protein degradation marker Atrogin mRNA content (Figure 8B, p=0.178). The global p-value for protein degradation marker MurF1 approached significance (Figure 8B, p=0.106). AN-5R rats had ∼300% greater Gadd45a mRNA content compared to CON-0R (p=0.005), but no other differences between groups were noted between either AN-0R or AN-15R (Figure 8B, p=0.954 and p=0.999 respectively). Autophagy marker Lc3 mRNA content was ∼67%, ∼51%, and ∼55% lower in AN-0R, AN-5R, and AN-15R compared to CON-0R, respectively (Figure 8B, p<0.001, p<0.001, and p<0.001 respectively). Additionally, another marker for autophagy, p62 mRNA content was ∼70%, ∼54%, and ∼70% lower in in AN-0R, AN-5R, and AN-15R compared to CON-0R, respectively (Figure 8B, p<0.001, p<0.001, and p<0.001 respectively). Following long-term recovery, there were no differences in Il6, Tnfα, or Myostatin mRNA content in AN-30R compared to CON-30R (Figure 8C, p=0.379, p=0.395 and p=0.253). In addition, there were no differences between AN-30R and CON-30R groups in Atrogin (p=0.423), MurF1 (p=0.810), Gadd45a (p=0.609), Lc3 (p=0.148), or p62 mRNA content (∼30% greater in AN-30R, p=0.08, Figure 8D).

**Figure 8:**
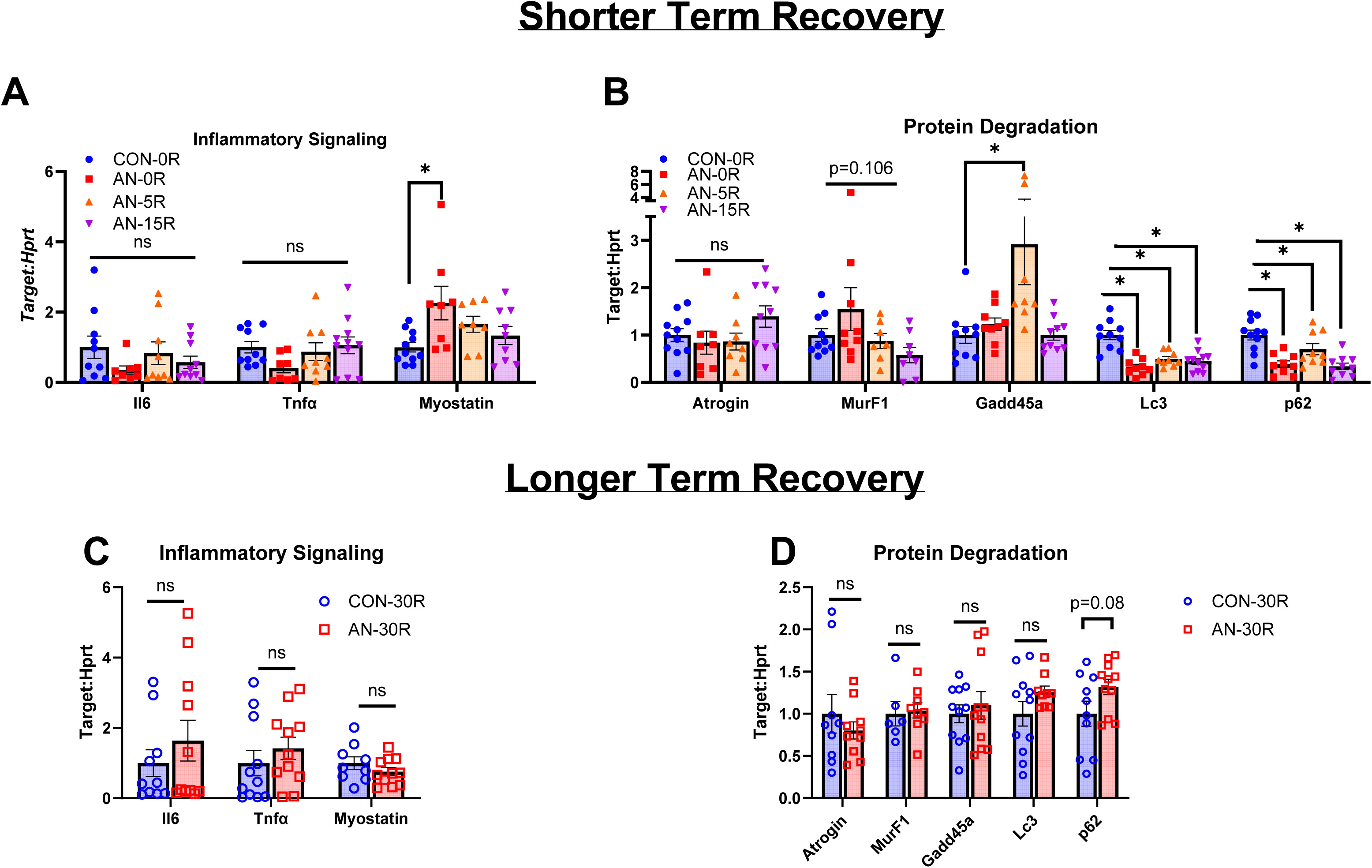
mRNA data for inflammatory signaling and protein degradation pathways. A) mRNA content for genes related to inflammatory signaling after simulated AN and shorter term recovery. B) mRNA content for genes related to proteasome ubiquitin and autophagy-mediated protein degradation signaling after simulated AN and shorter term recovery. C) mRNA content for genes related to inflammatory signaling following simulated AN and longer term recovery. D) mRNA content for genes related to proteasome ubiquitin and autophagy-mediated protein degradation following simulated AN and longer term recovery. CON-0R=healthy controls, AN-0R=simulated anorexia nervosa, AN-5R= simulated anorexia nervosa followed by 5 days of weight recovery, AN-15R= simulated anorexia nervosa followed by 15 days of weight recovery. CON-30R=healthy controls age-matched to AN-30R. AN-30R=simulated anorexia nervosa followed by 30 days of weight recovery. Data are depicted as mean ± SEM, *=p<0.05.

## Discussion

In our rodent model of simulated AN and subsequent recovery, we find multiple components of muscle size and strength are altered with AN, and many muscle features of AN are not resolved with weight recovery. Moreover, AN appears to induce changes in mRNA content of some moderators protein synthetic signaling, some of which persist following weight gain. In aggregate, these data confirm prior clinical reports of muscle weakness in individuals with AN^6^ and suggest muscle implications of AN may be more persistent than previously appreciated.

Overall, 30 days of simulated AN in rats (theoretically corresponding to roughly 2-3 years of AN in humans^24^) results in loss to both muscle size and strength. While many of these size alterations are restored to or even exceed levels in control rats with sufficient recovery, deteriorations in muscle strength appear particularly persistent. For example, AN-30R rats had lower volitional and nerve-stimulated force production (plantar flexion), yet there was a full restoration in muscle mass. This incongruity suggests alterations to inherent muscle quality due to previous AN. To our knowledge, most clinical studies of AN evaluate either changes in muscle/lean body mass, with few evaluating muscle strength^6^. In other myogenic pathologies associated with decrease in muscle quality, such as muscular dystrophy or volumetric muscle loss, this reduction in muscle quality is typically attributed to infiltration of non-contractile tissue such as fat^25, 26^ and/or collagen^27–29^. The (non-significant) lower muscle density we note in AN-30R rats may suggest some intramuscular fat accumulation. Conversely, prior studies in mice using a simulated model of relative energy deficiency in sports (RED-S), found significant alterations to collagen signaling during low-weight status^30^. Speculatively, the refeeding process may hyperactivate either fat or collagen deposition processes, allowing for a restoration of muscle size, without a corresponding restoration of muscle strength, resulting in lower overall muscle quality. At this point, we are unable to directly disentangle precise mechanisms involved in this reduction of muscle quality; however, our results suggest that simply assessing muscle size may not provide a comprehensive or accurate evaluation of muscle health in those with AN or a prior history of AN.

We note the musculoskeletal effects of AN appear to disproportionately affect muscles of different metabolic phenotypes and fiber-type characteristics. For example, we note significant and sustained changes in gastrocnemius function and size, whereas these alterations are less pronounced or persistent in the tibialis anterior and plantaris muscles. Moreover, alterations to muscle fiber CSA disproportionally affected MyHC I fibers as opposed to MyHC II fibers. In general, the gastrocnemius has a greater relative proportion of (presumably oxidative) MyHC I fibers compared to the tibialis anterior in the rat^31^. These data imply that AN, like many other atrophic conditions such as disuse atrophy and cancer cachexia^20–22^, has differential effects on different muscle phenotypes. AN appears to follow a similar pattern of muscle atrophy to disuse atrophy, whereby type I fibers appear particularly susceptible, compared to other fiber types^20, 32^. Correspondingly, these data suggest researchers and clinicians should apply discretion when evaluating specific muscles during AN and weight recovery. It is also noteworthy, for this study we performed most of our analysis on the gastrocnemius muscle — given the soleus’s high distribution of oxidative fibers^31^, it is plausible the soleus may have more physiological alterations during AN compared to the gastrocnemius. Indeed, a recent report in RED-S mice suggests a greater transcriptomic response to severe energy deficiency in the soleus compared to the gastrocnemis^30^. However, in this study we were only able to evaluate molecular signaling within the gastrocnemius muscle; correspondingly this speculation would need further validation in other muscles with different metabolic phenotypes. Finally, although speculative, given the known sex differences in fiber-type distribution between males and females, whereby females tend to have a greater proportion of oxidative muscle fibers^33^, it seems plausible that AN-induced muscle loss would affect females to a greater extent compared to males. Recent data in mice during RED-S seems to suggest a greater transcriptomic response in females relative to males^30^. However, this hypothesis would require further investigation to directly compare effects of energy deficiency and subsequent weight restoration between males and females.

In contrast to our original hypothesis, we found no changes in overall muscle protein synthesis between our AN and control or recovery rats. This contrasts with previous clinical research finding reductions in whole-body protein synthesis in patients with AN^9^. This discrepancy may in part be due to differences between whole-body vs. tissue specific protein synthesis. In previous studies using rodent models of AN, evaluation of colonic protein synthesis found reductions in protein synthesis as well as in corresponding mechanisms associated with protein synthesis^34, 35^. Moreover, duration of starvation or energy deficiency may play a large role in the magnitude of changes in protein synthetic rate. In prior studies of pure starvation as well as semi-starvation, dramatic changes to protein synthesis and degradative pathways are evident within the first 24-48 hours^36–38^. Whereas given sufficient time of starvation (over the course of multiple days/weeks), protein synthetic and degradative pathways appear to “normalize” ^37^. Based on these previous studies, it seems plausible that AN may initially result in dramatic changes to protein turnover pathways, which subsequently reach a new equilibrium given sufficient time, despite sustained muscle atrophy during AN. However, previous studies on this subject focused primarily on short-term dramatic starvation designs, as opposed to a more gradual weight loss which is more typical of clinical AN. Therefore, more research is necessary to evaluate how these protein synthetic pathways are altered during the development of AN and how these effects relate to changes in muscle size and strength. It is noteworthy that our AN-30R rats sustained decreases in Igf1 mRNA content and elevations in Deptor and Redd1 mRNA content. Igf1 facilitates protein synthesis, whereas both Deptor and Redd1 inhibit protein synthesis via mTOR repression^39, 40^. Taken together, sustained reductions in Igf1 in combination with increased Redd1 and Deptor (particularly in the context of the fed state) would suggest a less favorable acute protein synthetic response with previous AN. These cellular alterations would not necessarily be associated with differences in the 24-hour protein synthesis rate, but may suggest long-term changes to anabolic responses that could have long-term effects on musculoskeletal health.

There are a few limitations of our present study that should be acknowledged. While we find many components of muscle size and strength are normalized in AN-15R rats, we must acknowledge these rats are 15 days older than CON-0R rats. Due to the continued maturation of rats at this age, it is possible that AN-15R rats return to a baseline of muscle size/strength, but may still have lower muscle strength/size compared to age matched controls. Given our findings in AN-30R and CON-30R rats, this hypothesis seems plausible and may need further exploration. Additionally, our molecular data were collected in a matched fed state; it is possible some of the observed differences (or lack thereof) would be different in a completely fasted state or with a naturally occurring feeding pattern (i.e. Control/AN-recovery rats with fully ad libitum feeding through the night and AN rats fasted), this may warrant further investigation in the future. Moreover, mRNA content represents only one component of overall cellular signaling and does not fully capture the interactions of processes such as DNA methylation/epigenetic alterations, ribosomal translation, or protein phosphorylation and functionality, which would need to be explored in future studies. In addition, while we did extend our evaluation to 30 days of recovery, additional studies assessing animals for several months would be valuable to better understand even longer-term sequalae. Finally, although we believe our model captures the physiological features of weight recovery following AN, full recovery from AN is a complex process and rarely is as simple as mere weight gain. There are often relapses^41^ with corresponding fluctuations in bodyweight and presumably muscle health. Similarly, it is unclear if the physiological and molecular responses we have noted in this study would directly mirror the clinical condition over prolonged recovery periods.

In conclusion, in our rodent model of AN and weight restoration, we find significant alterations in muscle size and strength with AN. Given sufficient time, muscle size recovers to healthy control levels; however, there appear to be longer-term alterations to muscle strength, suggesting a decrease in overall muscle quality following AN. These changes correspond to changes in protein synthetic signaling; for example, anabolic signaling cascades appear attenuated following long-term recovery from AN. These results suggest that musculoskeletal implications of AN may be longer-lasting than previously appreciated and may be a key contributor to increased morbidity in those ongoing or previous AN.

## Data Availability Statement

Raw data and associated statistical code are available at our Open Science Framework page for this project.

## Acknowledgements

This study was supported by NIH grant K12HD051959 (M.E.R). We would like to thank Eliza Garland, Sheridyn Pinkham, Madisyn Hancock, and Sophie Jalkut for their assistance with this project.

